# Genetic Association Studies in Transgender Cohorts: A Systematic Review and Meta-Analysis

**DOI:** 10.1101/2023.02.27.530343

**Authors:** Blake Ashley, Vincent Harley

## Abstract

According to twin studies, there is a heritable contribution to gender incongruence, but the genetic mechanisms of this are unknown. Recent efforts to identify an aetiology of gender incongruence have focused on the hypothesis that sex hormones establish gender identity through influencing the development of neuroanatomy. Candidate gene studies that have sought to elucidate whether polymorphisms in sex steroidogenesis genes are overrepresented in transgender populations have been equivocal. A systematic search for case-control genetic association studies in transgender populations was conducted. Mean (+SD) or allele frequencies were extracted and combined quantitatively in random effects meta-analysis, summarised as standardised mean difference for continuous alleles or odds ratios for allele frequencies. Eight studies were included in the analysis. These studies spanned polymorphisms in five genes; the CAG repeat in androgen receptor (*AR*), the TA repeat in estrogen receptor 1 (*ESR1*), the CA repeat in estrogen receptor 2 (*ESR2*), the TTTA repeat in cytochrome P450 family 19 subfamily A member 1 (*CYP19*), and the T>C SNP in cytochrome P450 family 17 subfamily A member 1 (*CYP17*). Pooled estimates indicated that transgender women have a significant overrepresentation of short *ESR1* alleles compared to cisgender men (OR = 1.23, 95% CI: 1.06, 1.44, p = 0.0089). This may contribute an increased likelihood of developing gender incongruence amongst natal males. Future investigations into gender incongruence should use genome-wide methods.

## Introduction

Gender identity refers to one’s psychological experience of their own gender. For most, physical sex and gender identity are congruent. However, for some individuals the two can be incongruent; their gender identity does not match their physical sexual characteristics. This incongruence can be distressing, and at clinically relevant levels, can be diagnosed as gender dysphoria according to the Diagnostic Statistical Manual of Mental Disorders (DSM-5) (American Psychiatric Association, 2013), or gender incongruence (GI) according to the International Classification of Diseases (ICD-11) (World Health Organisation, 2022). GI will be used throughout this paper to refer to the state of having an incongruence between gender identity and biological sex. Transgender men are individuals who were born physically as female but identify as men. Transgender women are individuals who were born physically as male but identify as women.

Recent conceptions of an aetiology of GI have incorporated biological factors. The prevailing hypothesis is that gender identity is established during foetal/early post-natal development via sex hormones influencing the development of the brain. The hypothesis supposes that in transgender individuals, the process by which sex hormones influence the development of gender identity is altered (Guillamon et al., 2016). This idea has its basis in the organisational/activational dogma of hormonal impacts on the sexual differentiation of the brain. Under this framework, sex hormones prenatally organise brain tissues through receptor interactions, which later manifests in sex-specific behaviours when ‘activated’ by subsequent hormonal release (Arnold, 2017). Given that gender identity is one of few variables strongly correlated with biological sex (i.e., most biological males also have a male gender identity), the current hypothesis of biological contributions to gender identity posits that it may be a sex-typical behaviour that is, at least in part, prenatally instantiated by the influence of sex hormones. Evidence for this hypothesis primarily stems from studies in people with Differences of Sex Development (DSDs), where variants in sex steroidogenesis genes alter the development of chromosomal or anatomical sex. DSDs are associated with increased rates of cross-gender identification and gender incongruence, such as in congenital adrenal hyperplasia (Dessens et al., 2005; Pasterski et al., 2015; Seneviratne et al., 2021), partial androgen insensitivity syndrome (Mazur, 2005), and 5-alpha reductase type 2 deficiency (Cohen-Kettenis, 2005). Neuroanatomical studies have also demonstrated that transgender individuals displays differences to cisgender individuals in various brain regions (Mueller et al., 2021), however the degree to which these findings support aetiological hypotheses or whether they are merely neurobiological representations of the experience of gender incongruence is up for debate.

Though direct evidence for genetic variants functionally affecting sex hormone signalling in transgender participants is limited, twin data does suggest a substantial genetic component, with concordance rates in monozygotic twins being greater (37%) than in dizygotic twins (0%) (Heylens et al., 2012). Most genetic-oriented study in transgender populations have been candidate gene studies comparing the frequency of polymorphisms in various sex hormone signalling genes between cisgender and transgender participants. The selection of these specific genes has been based on the hypothesis outlined above. These polymorphisms are thought to be functional, given their associations to other phenotypes. For example, the trinucleotide CAG repeat polymorphism in exon 1 of the androgen receptor gene (*AR*(CAG)n), the dinucleotide CA repeat in intron 5 of the estrogen receptor 2 gene (*ESR2*(CA)n), the TTTA repeat in intron 4 of cytochrome P450 family 19 subfamily A member 1 (*CYP19*(TTTA)n), and the T>C SNP in the 5’ UTR of the cytochrome P450 family 17 subfamily A member 1 (*CYP17*) gene have been associated with hormone levels in adulthood (Gennari et al., 2004, 2005; Scariano et al., 2004; Sharp et al., 2004; Westberg et al., 2001). As well as the dinucleotide TA repeat in the promotor region of the estrogen receptor 1 gene (*ESR1*(TA)n), these polymorphisms are also associated with fertility (Xiao et al., 2016), prostate cancer risk (Weng et al., 2017), and bone mineral density (Gennari et al., 2005; Langdahl et al., 2000; Ogawa et al., 2000). Further, *AR*(CAG)n correlates with male-typical patterns of brain structure (Perrin et al., 2008; Raznahan et al., 2010). *ESR1*(TA)n has also been associated with the ratio of the length of the second finger to the fourth finger (2D:4D) (Vaillancourt et al., 2012), which shows sex differences (Hönekopp & Watson, 2010) and is thought to be a proxy measure of prenatal hormone exposure (McIntyre, 2006). This is especially relevant in the context of GI, as transgender women show feminised 2D:4D ratios compared to controls (Siegmann et al., 2020).

The results of candidate gene studies in transgender cohorts have often been contradictory. Longer *AR*(CAG)n have been found more frequently in transgender women (Hare et al., 2009), though this has failed to replicate in several other studies (Fernández, Esteva, Gómez-Gil, et al., 2014; Fernández et al., 2018; Foreman et al., 2019; Henningsson et al., 2005; Lombardo et al., 2013; Ujike et al., 2009). However, three studies have found that *AR*(CAG)n was significantly associated with transgender women when in conjunction with alleles of other sex steroidogenesis genes, including *CYP19* (Henningsson et al., 2005), *PGT* and *COMT* (Foreman et al., 2019) and *ESR2* (Fernández et al., 2018; Foreman et al., 2019). Notably, a previous meta-analysis comparing mean *AR*(CAG)n in transgender women and cisgender men found that transgender women had significantly, though very slightly, longer repeat lengths (D’Andrea et al., 2020). However, the analysis did not include data from the second largest association study in transgender women (Foreman et al., 2019), and modelling was used to estimate mean *AR*(CAG)n length in one of the studies (Fernández et al., 2018), even though exact allele frequencies were available in the supplementary data, potentially causing inaccuracies. Two studies examining *AR*(CAG)n in transgender men found no significant relationships (Fernández et al., 2018; Ujike et al., 2009). Shorter *ESR1*(TA)n were found in transgender women in one study (Foreman et al., 2019) but not others (Cortés-Cortés et al., 2017; Fernández, Delgado-Zayas, et al., 2020; Ujike et al., 2009), whilst transgender men showed no difference in *ESR1*(TA)n compared to cisgender women in three studies (Cortés-Cortés et al., 2017; Fernández, Delgado-Zayas, et al., 2020; Ujike et al., 2009). CA repeat length in *ESR2* has been associated to both transgender women (Henningsson et al., 2005) and transgender men (Fernández et al., 2018), though these results have also failed to replicate in other studies (Fernández et al., 2018; Foreman et al., 2019; Hare et al., 2009; Ujike et al., 2009). No studies have demonstrated an independent association between *CYP19* and gender incongruence in transgender women or transgender men (Fernández, Esteva, Gómez-Gil, et al., 2014; Fernández, Esteva, Gómez Gil, et al., 2014; Fernández et al., 2018; Foreman et al., 2019; Hare et al., 2009; Ujike et al., 2009). Only one study has shown an association between *CYP17* and gender incongruence (Bentz et al., 2008), whilst others have not (Fernández et al., 2015; Foreman et al., 2019).

The equivocal results of these studies are likely due to small sample sizes and study heterogeneity. For example, the results of genetic association studies are contingent on the ethnicity of the participants, which may differ between studies. Furthermore, transgender individuals, even within the same natal sex, may not be a homogenous group with the same developmental trajectories. A conceptual distinction has been made between transgender individuals who had early onset of GI and tend to be attracted to members of their natal sex, whilst transgender individuals who have later onset tend to be attracted to the sex of their gender identities (Blanchard, 1985; Guillamon et al., 2016). Thus, studies may differ in their results due to differences in the subtypes of transgender participants recruited. Inconsistency between study results has also contributed to disagreement in the literature about the degree to which candidate gene studies support the hypothesis that GI has a basis in altered sex hormone signalling. For example, some cite genetic association studies as supporting the hypothesis that GI has at least a partial basis in altered sex hormone signalling during early development (Polderman et al., 2018; Rosenthal, 2021), whilst some claim that results are too nascent and contradictory to offer such evidence (Levin et al., 2022).

To gain further insight into the role, if any, of sex steroidogenesis genes in a genetic contribution to GI, we conducted a systematic review and meta-analysis of previous candidate gene studies. In studying the mechanisms by which GI may occur, we also investigate how gender identity develops in general. We hypothesised that specific polymorphisms in genes related to sex hormone signalling would be overrepresented in transgender populations compared to cisgender populations, and thus that sex steroidogenesis genes play a role in the development of gender identity.

## Methods and Materials

This study was conducted in accordance with the Preferred Reporting Items for Systematic Reviews and Meta-Analysis guidelines (Page et al., 2021) and the Meta-Analysis of Observational Studies in Epidemiology checklist (Brooke et al., 2021).

### Eligibility Criteria

Eligibility for inclusion in this analysis were: a) case-control genetic association study, b) comparison of allele length/frequency between transgender women (cases) and cisgender men (controls), or between transgender men (cases) and cisgender women (control), and c) availability of either mean length and standard deviation (SD), or allele frequencies, or adequate information to derive these, for both cisgender and transgender cohorts. When participants were included in multiple studies, the study with the largest sample size was included.

### Search Strategy

MedLine, Scopus, and the Cochrane Library databases were searched. Searches were done for papers published between January 2000 and August 2022. No restriction was put on the language of the publication. The database search strategy was designed and conducted by B.A, with guidance from Monash Health librarians on incorporating MeSH terms. In all three databases, a range of keywords were used; “transgender”, “gender incongruence”, “gender dysphoria”, “gender identity disorder”, “transsex*”, “MTF”, “FTM”, “assigned sex”, “trans?women”, “trans?men”, “transgender women”, “transgender men”, “polymorphism*”, “variant*”, “allele*”, “repeat”, “SNP”, “RFLP”, “sex steroid*”, “sex hormone*”, “hormone*”, “receptor*”, “androgen receptor*”, “estrogen receptor” “oestrogen receptor*”, “metabolism”, “enzyme”, “androgen*”, “testosterone”, “estrogen”, “oestrogen”, “aromatase”, “CYP17”. In the MedLine and Cochrane Library databases, MeSH terms were also used; “Transgender Persons”, “Gender Dysphoria”, “Transsexualism”, “Polymorphism, Genetic”, “Polymorphism, Restriction Fragment Length”, “Polymorphism, Single Nucleotide”, “Alleles”, “Gonadal Steroid Hormones”, “Testosterone”, “Estrogens”, “Aromatase”. Reference lists of retrieved articles were also scrutinised to identify further potential studies, but no additional studies were found. Once the search was complete, titles and abstracts were retrieved and reviewed for relevance. The full texts of publications that passed this stage were then read to deem eligibility.

### Data Extraction

Data was extracted by both B.A and V.H. From the eligible studies, first author, year, geographic location, age of participants, diagnostic tool used to define cases, the gene(s) examined, and allelic information for meta-analysis were extracted. In some cases, studies analysed polymorphisms in different manners and thus reported different statistical endpoints. For example, differences in *AR*(CAG)n were analysed by both t-tests (Henningsson et al., 2005; Lombardo et al., 2013), Mann Whitney tests (Fernández et al., 2018; Foreman et al., 2019; Ujike et al., 2009), and through dichotomisation of the repeat into short and long forms and subsequent calculation of odds ratios (Henningsson et al., 2005). Thus, before data extraction took place, it was decided that a) normally distributed repeat length fragment polymorphisms (RFLPs) would be meta-analysed via the calculation of a pooled standardised mean difference estimate, b) non-normally distributed RFLPs would be meta-analysed via calculation of a pooled odds ratio estimate, and c) single nucleotide polymorphisms would be meta-analysed via calculation of a pooled odds ratio estimate. For normally distributed RFLPs, it was therefore necessary to obtain the means and standard deviations for both case and controls groups. In cases where these were not explicitly reported, there were calculated using a method that estimates mean and standard deviation from distribution information (Wan et al., 2014), or obtained via contact with the original authors. For non-normally distributed RFLPs, dichotomised allele frequencies, where short alleles were defined as equal to or less than the median allele length in the corresponding control group, were either obtained directly from publications or via contact of the original authors or calculated based on allele distributions supplied in the publications. Inconsistency in data extraction were resolved via discussion and re-extraction of the data to reach consensus.

### Quality of Studies Assessment

The Newcastle-Ottawa Scale (NOS) (Mamikutty et al., 2021; Wells et al., 2014) was used to assess the quality and risk of bias of the included studies. This scale measures study quality along three domains; selection of participants, comparability of groups, and the measurement of outcome. Studies can receive a maximum of 9 points and a minimum of 0. Scores of 0-4 are deemed low quality, 5 to 6 moderate, and 7-9 high quality. Study quality was assessed by both B.A and V.H. Consensus was reached through discussion after assessment was conducted individually. Inter-rater reliability was assessed via calculation of Cohen’s kappa.

### Statistical Analysis

The association between *AR*(CAG)n and GI was assessed via calculation of the pooled standardised mean difference (SMD) with a 95% confidence interval. The associations between *ESR1*(TA)n, *ESR2*(CA)n, *CYP19*(TTTA)n, and *CYP17* (T>C) were assessed via calculation of a pooled odds ratios with 95% confidence intervals, quantifying the strength of the association between form of the allele (short/long for *ESR1*, *ESR2* and *CYP19* and T/C for *CYP17*) and GI. Heterogeneity between the studies was assessed using the I^2^ test and the Cochrane Chi-square *X*^2^ (Cochrane Q) statistic, where heterogeneity was defined as I^2^ = >50%, and p = <0.1, respectively (Cordero & Dans, 2021; Petitti, 2001; Song et al., 2001). In cases of significant heterogeneity, leave-one-out sensitivity analysis was conducted to assess which studied contributed significant heterogeneity. This was only carried out in analyses that contained >3 studies. Given the studies included participants from different ethnic backgrounds, random effects models were used to calculate pooled effect estimates in all instances. The results of fixed and random effects models are essentially identical in the absence of heterogeneity (Petitti, 2001), but random effects models are necessitated in instances where there may be systematic differences in study characteristics, such as in the ethnicity of the participants (Munafò et al., 2004).

Publication bias was assessed through visual examination of funnel plots and statistically with the use of a regression test (Egger et al., 1997), followed by the trim-and-fill method (Duval & Tweedie, 2000) to attempt correction of any suspected publication bias. Given that the sensitivity of statistical methods in assessing publication bias is low for meta-analysis with few studies (Munafò et al., 2004), publication bias was only tested for in this study where analysis involved ≥ 3 studies, and p < 0.1 was used as the significance cut off for the regression test to account for the lack of power. All analysis and visualisation was conducted using R Studio (v. 1.4.1717) with the ‘metafor’ package (v3.0.2) (Viechtbauer, 2010).

## Results

### Study Selection

After database searches and screening for inclusion/exclusion criteria, eight studies were included in the quantitative analysis (figure 1). Amongst these studies, polymorphisms in 5 genes were examined in multiple (>2) studies, presenting the opportunity for meta-analysis. These polymorphisms were *AR*(CAG)n, *ESR1*(TA)n, *ESR2*(CA)n, *CYP19*(TTTA)n, and *CYP17* (T>C).

**Figure 1:**
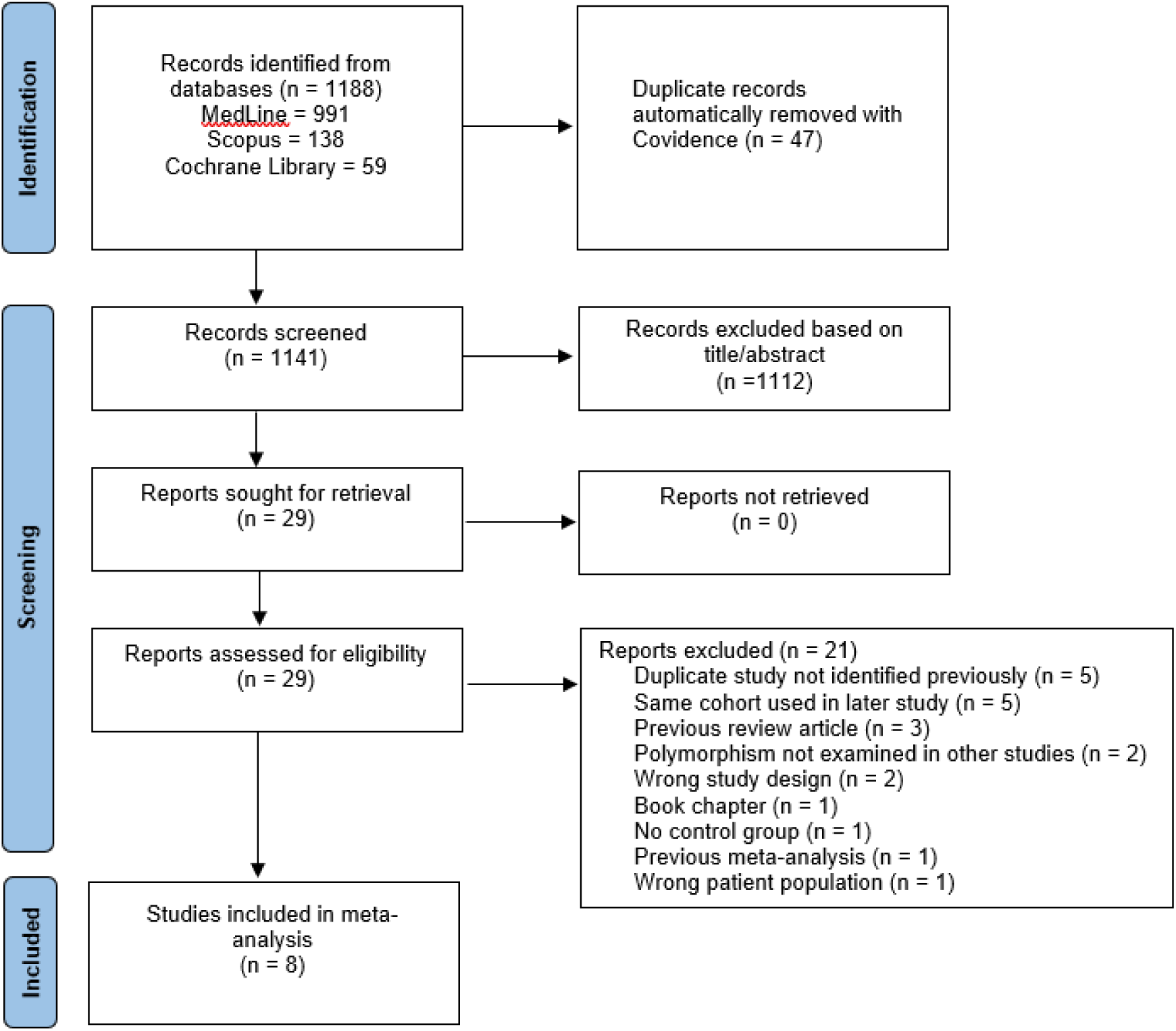
PRISMA flow chart for the study selection process.

### Study Characteristics

**Table 1:**
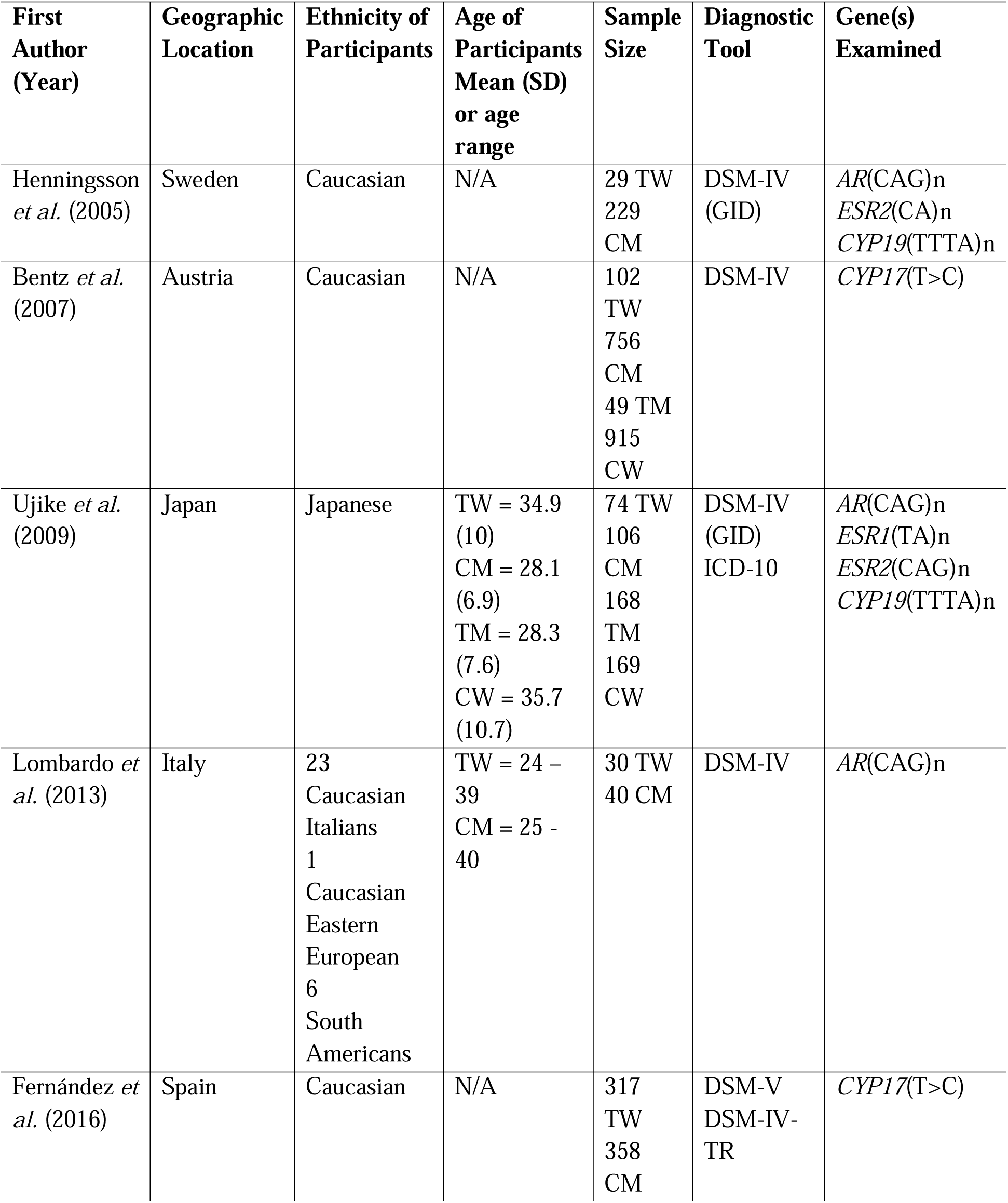
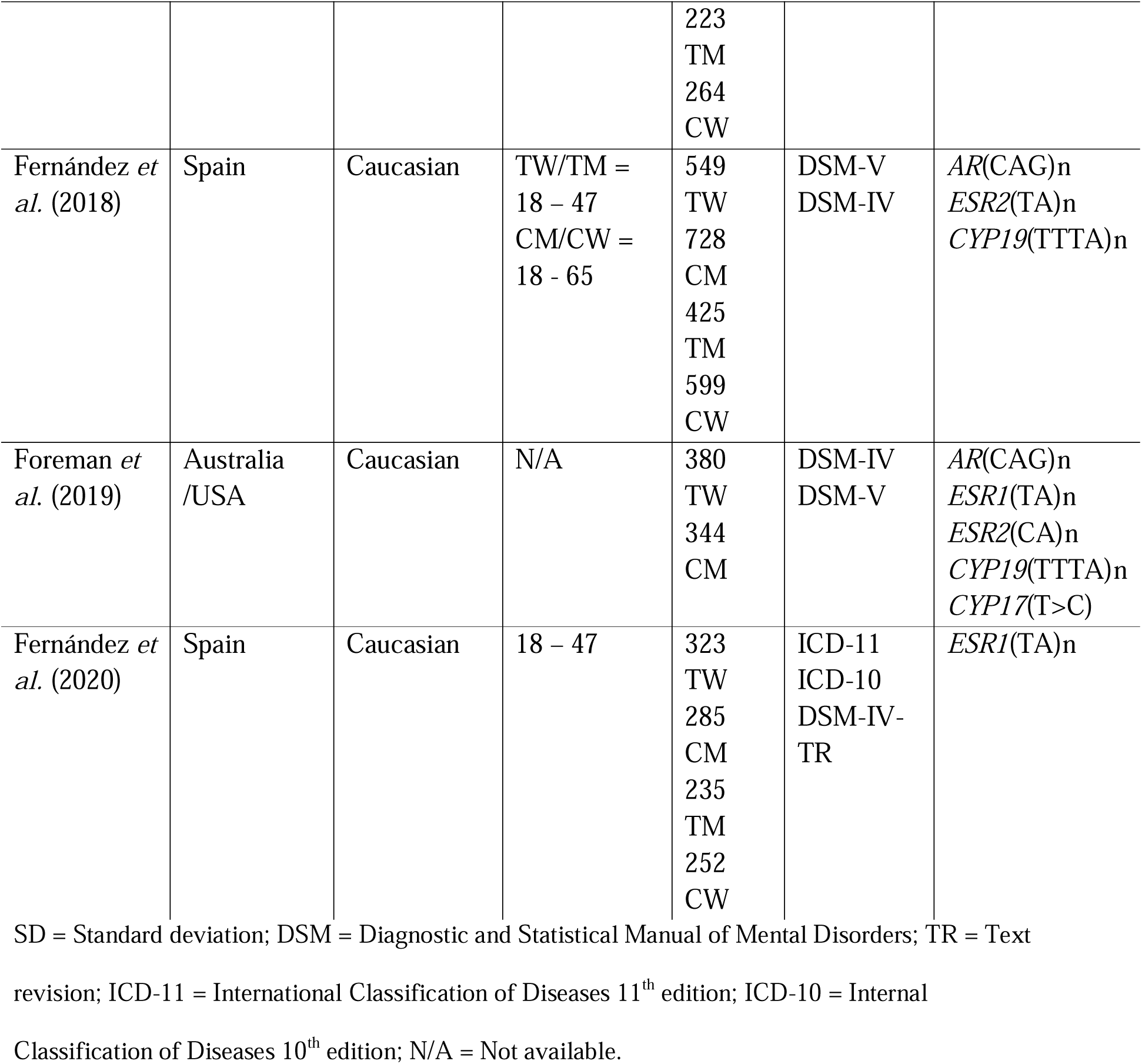
Characteristics of studies included in meta-analysis.

### Quality of Included Studies

The quality ratings of studies included in the meta-analyses are shown in table 1. Three of the studies were rated low quality, two were rated moderate quality, and two were rated high quality. Inter-rater reliability was substantial (k = 0.76) before consensus.

**Table 2:** Newcastle-Ottawa Assessment Scale assessment of included studies. Total scores of 0-4 are deemed low quality, 5 to 6 moderate, and 7-9 high quality.

| Study (First Author) | Selection |  |  |  | Comparability |  | Outcome |  |  | Total |
| --- | --- | --- | --- | --- | --- | --- | --- | --- | --- | --- |
|  | Definition of Cases | Representativeness of Cases | Selection of Controls | Definition of Controls | Controls for Ethnicity | Controls for Other Factors | Assessment of Exposure | Same Ascertainment Method of Cases/Controls | Non-Response Rate |  |
| Henningsson *et al.* (Henningsson *et al.*, 2005) | 1 | 0 | 1 | 0 | 1 | 0 | 1 | 1 | 0 | 5 |
| Bentz *et al.* (Bentz *et al.*, 2008) | 1 | 0 | 0 | 0 | 1 | 0 | 1 | 1 | 0 | 4 |
| Ujike *et al.* (Ujike *et al.*, 2009) | 1 | 0 | 0 | 1 | 1 | 0 | 1 | 1 | 0 | 5 |
| Lombardo *et al.* (Lombardo *et al.*, 2013) | 1 | 0 | 0 | 1 | 0 | 0 | 1 | 1 | 0 | 4 |
| Fernández *et al.* (Fernández *et al.*, 2016) | 1 | 0 | 1 | 1 | 1 | 1 | 1 | 1 | 0 | 7 |
| Fernández *et al.* (Fernández *et al.*, 2018) | 1 | 0 | 1 | 1 | 1 | 1 | 1 | 1 | 0 | 7 |
| Foreman *et al.* (Foreman *et al.*, 2019) | 1 | 0 | 0 | 1 | 0 | 0 | 1 | 1 | 0 | 4 |
| Fernández *et al.* (Fernández, Delgado-Zayas, *et al.*, 2020) | 1 | 0 | 0 | 0 | 1 | 1 | 1 | 1 | 0 | 5 |

### Synthesis of Association Studies

Figures 2 and 3 display the results of the meta-analyses in transgender women/cisgender men, and transgender men/cisgender women, respectively. Out of all analyses, one significant association was identified; transgender women had a significant overrepresentation of short *ESR1*(TA)n compared to cisgender men (p = 0.0089) (Figure 2a). Heterogeneity tests found significant heterogeneity between the results of the individual studies in the analysis of *ESR2* in transgender women (figure 2a). Subsequent sensitivity analysis revealed that all heterogeneity disappeared when the Henningsson et al., (2005) study, which had a much smaller sample size than the other studies, was removed. After removal, the model became more insignificant, with the OR tightening around 1 (OR = 0.98, 95% CI = [0.86, 1.12], p = 0.86). Significant heterogeneity was also identified between the studies in the analysis of *CYP19* in transgender women (figure 2a). Removal of Fernández et al. (2018) eliminated heterogeneity, but the model remained insignificant (OR = 1.19, 95% CI = 0.99, 1.43], p = 0.07).

**Figure 2:**
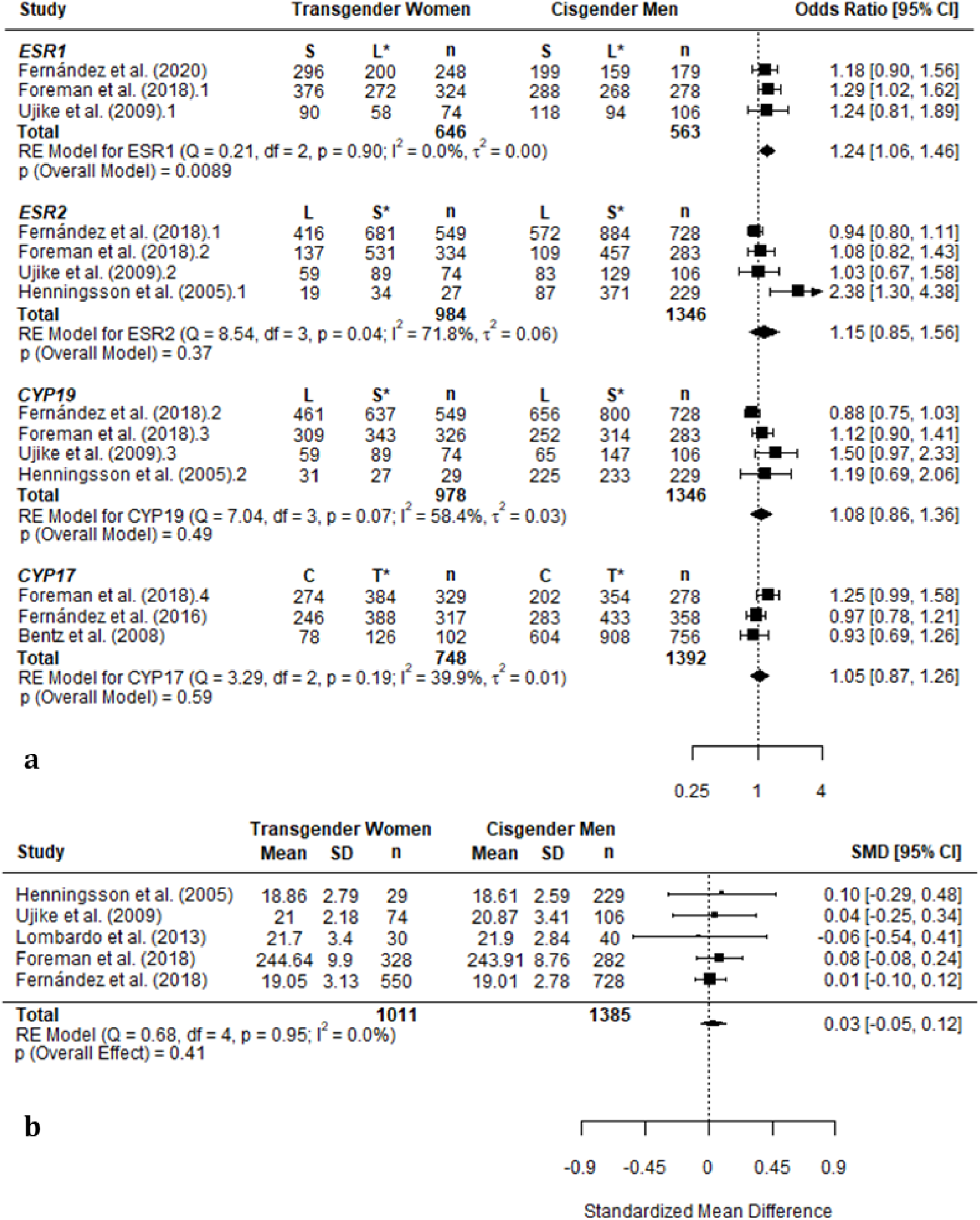
Forest plots of meta-analysis of dichotomised *ESR1*(TA)n, *ESR2*(CA)n, *CYP19*(TTTA)n and *CYP17* (T>C) (a) and mean *AR*(CAG)n (b) between transgender women and cisgender men. Diamonds represent overall summary estimates. Width of the diamond represents the 95% confidence interval. Boxes indicate the effect estimate obtained in the individual studies, and box width indicates the weight given to each individual study based on its sample size. Whiskers indicate 95% confidence intervals. Odds ratios are in reference to the reference allele for each gene, indicated by *. *ESR1*, *ESR2*, *CYP19*, and *CYP17* are autosomal genes, so the number of alleles is double the number of participants. For *AR*(CAG), Foreman *et al*. (2018) recorded *AR*(CAG)n as total length of the repeat sequence, rather than the number of trinucleotide repeats as other studies did. SD = standard deviation, n = number of participants, SMD = standardised mean difference, CI = confidence interval, RE = random effects, df = degrees of freedom

**Figure 3:**
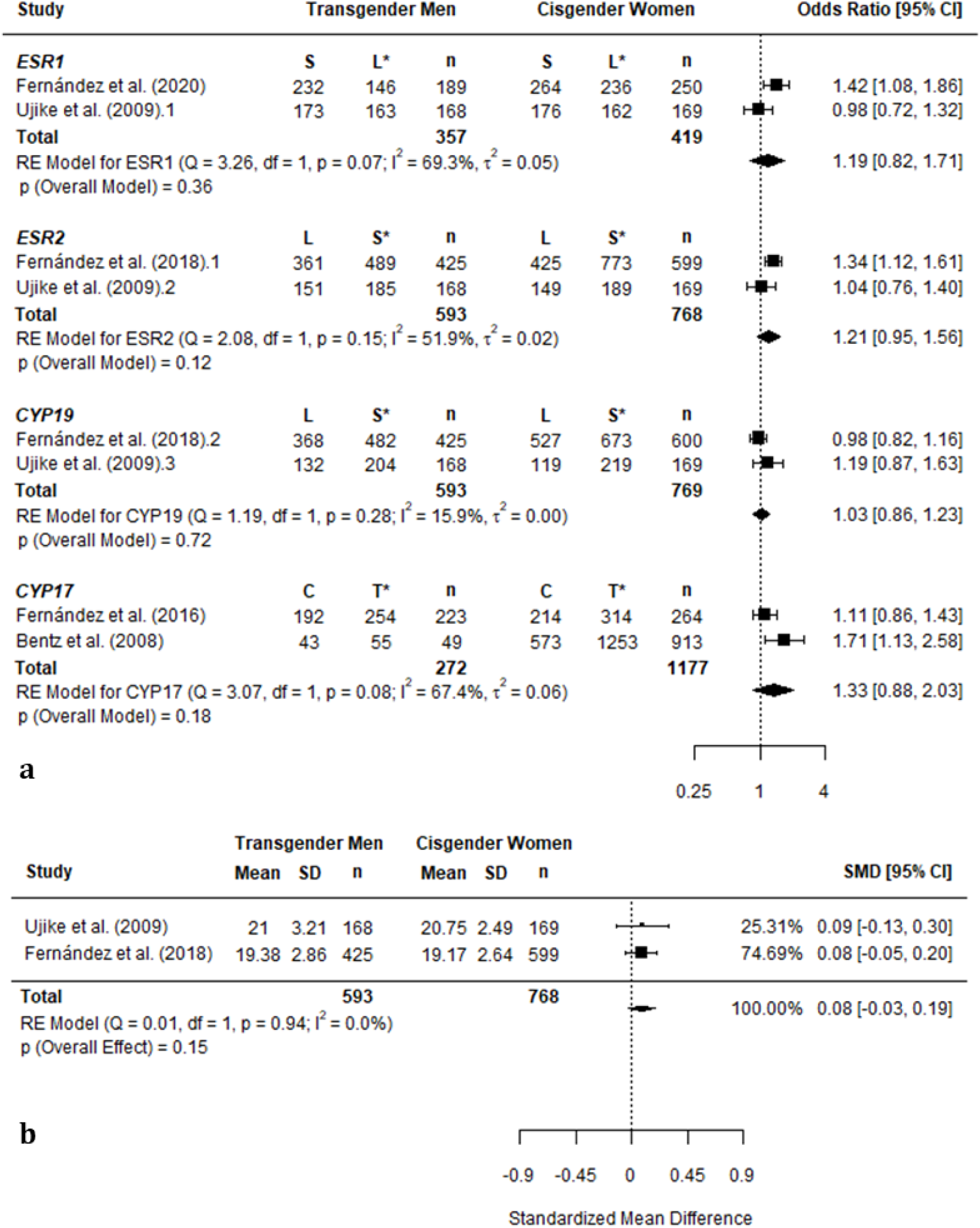
Forest plots of meta-analysis of dichotomised *ESR1*(TA)n, *ESR2*(CA)n, *CYP19*(TTTA)n and *CYP17* (T>C) (a) and mean *AR*(CAG)n (b) between transgender women and cisgender men.

### Publication Bias

Figure 4 displays the funnel plots obtained from testing for publication bias in the analyses of all five genes in transgender women and cisgender men. Publication bias was not tested for in the studies of transgender men as there were too few studies. Across *AR*, *ESR1*, *ESR2*, *CYP19*, and *CYP17*, regression tests for funnel plot asymmetry were not significant (p = 0.86, 0.76, 0.16, 0.17, 0.77, respectively). Trim-and-fill analysis did not result in any potentially missing studies being added to the funnel plots for *AR*(CAG)n, *ESR2*(CA)n, or *CYP17* T>C. Two putative missing studies were added to the right side of the funnel plot for *ESR1*(TA)n, and two putative missing studies were filled in on the left side of the plot for *CYP19*(TTTA)n. The model remained non-significant for *CYP19*(TTTA) after these putative studies were added (OR = 0.98, 95% CI = [0.80, 1.22], p = 0.88). For *ESR1*(TA)n, the model remained significant after these studies were added (OR = 1.28, 95% CI = [1.13, 1.46], p = 0.0002. Thus, no changes to the overall results from these meta-analyses occurred after testing for publication bias.

**Figure 4:**
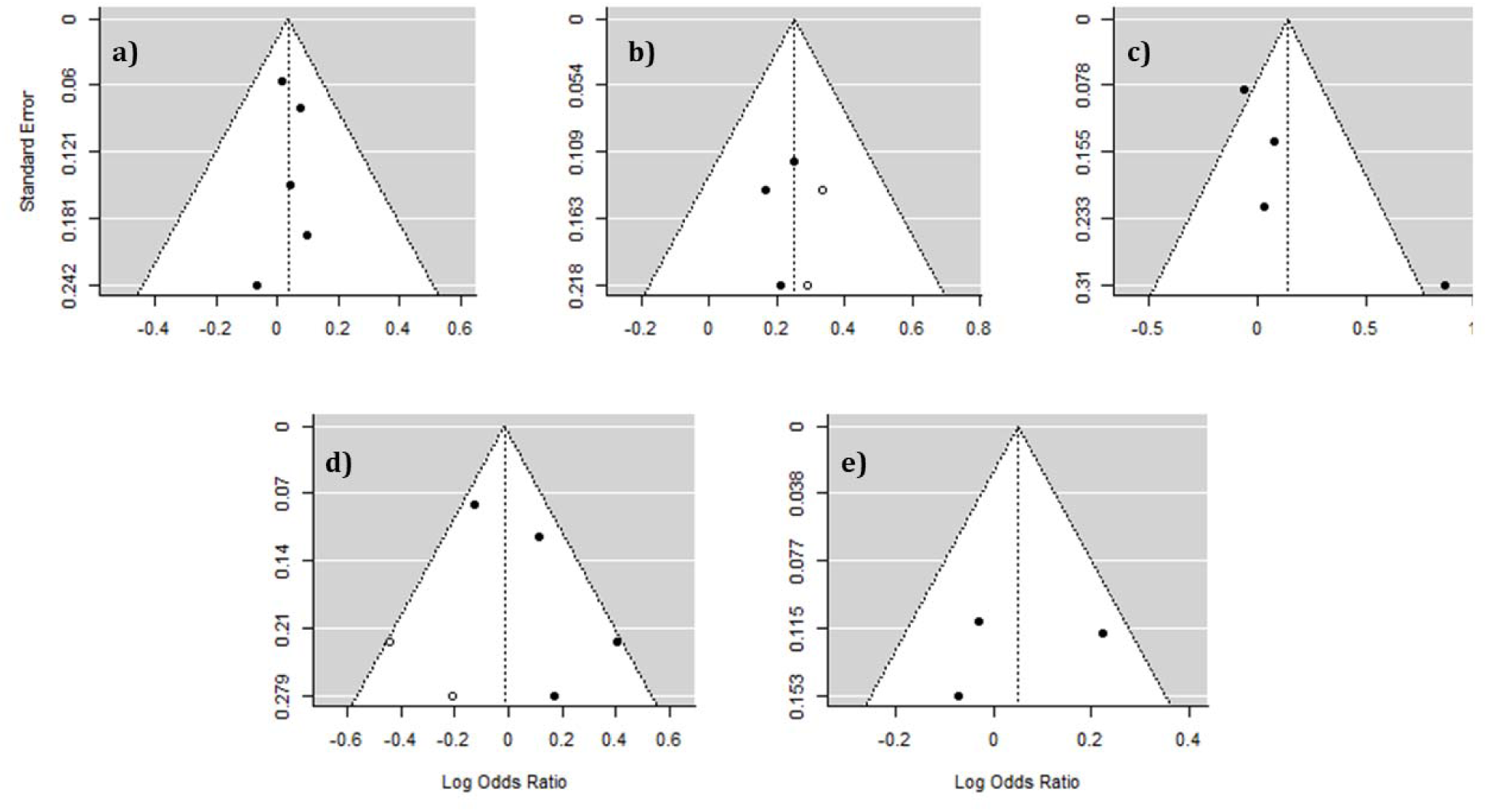
Funnel plots of meta-analyses of allele difference in *AR*(CAG)n (a), *ESR1*(TA)n (b), *ESR2*(CA)n (c), *CYP19*(TTTA)n (d) and *CYP19* T>C (e) in transgender women and cisgender men.

## Discussion

This is the largest genetic analysis of transgender individuals conducted to date. Mean *AR*(CAG)n, frequency of dichotomised short/long *ESR1*(TA)n, *ESR2*(CA)n, *CYP19*(TTTA)n and T>C SNP frequency in *CYP17* were compared between transgender women and cisgender men, and transgender men and cisgender women. One significant association was identified, where transgender women had a significantly higher frequency of short *ESR1*(TA)n, supporting our earlier finding (Foreman et al., 2019).

*ESR1* is highly expressed throughout the human brain (Laflamme et al., 1998). The length of the TA repeat in the promotor region of *ESR1* (*ESR1*(TA)n) correlates with the volume and density of many structures in the brain which display sex differences (Tan et al., 2020). However, there has been considerable debate regarding the role of estrogen receptors in the sexual differentiation of the brain. In rodents, masculinisation is typically accepted as occurring through the binding of locally aromatised testosterone to estrogen receptors, which does not occur in females due to the protective role of α-fetoprotein (Zuloaga et al., 2008). The role of estrogen receptors in human brain sexual differentiation is less clear. It has been suggested sexual brain differentiation in humans occurs primarily through direct binding of testosterone to androgen receptors (Puts & Motta-Mena, 2018), with female brain differentiation occurring merely due to the absence of testosterone. However, some argue that female brain differentiation is a more active process than this (Nugent et al., 2015), which involving the action of *ER*s (Bakker & Brock, 2010; Luoto & Rantala, 2018). If GI in natal males is the result of under masculinisation/feminisation of brain structures, our results may suggest that *ESR1* plays an active role in sexual brain differentiation.

The finding that short *ESR1*(TA)n are overrepresented in transgender women may support the idea that GI has its basis in altered sex hormone signalling influencing the development of sexually differentiated brain structures. However, it is currently difficult to tie in genetics findings with results from neuroanatomical studies in transgender individuals. For example, a diffusion tensor imaging study demonstrated the superior longitudinal fascicles are feminised in transgender women (Rametti et al., 2011). Feminisation of the region is correlated with short *ESR1*(TA)n (Tan et al., 2020), which seems in congruence with our results. However, in other brain regions that are feminised in transgender women, such as the hypothalamus (Garcia-Falgueras & Swaab, 2008; Zhou et al., 1995), short *ESR1*(TA)n is associated with masculinisation (Tan et al., 2020). Therefore, whilst estrogen signalling affects brain sexual differentiation, our ability to relate it to specific brain regions in transgender individuals is limited.

Other types of genetic analysis of transgender individuals have also implicated estrogen signalling. First, a whole-exome analysis identified 5 variants in estrogen signalling pathway genes that were unique amongst transgender individuals (Theisen et al., 2019), though this study only included 30 transgender individuals. Second, a candidate gene study of *SRC* genes showed certain SNPs were overrepresented amongst transgender individuals (Ramírez et al., 2021). SRCs are coactivators of estrogen receptors (Yore et al., 2010) and are influential in the masculinisation of the brain (Auger et al., 2000). Third, an epigenetic study (Fernández, Ramírez, et al., 2020) demonstrated sex differences in the degree of methylation in a regulatory region *ESR1*, and that both transgender women and transgender men had significantly lower levels of methylation in this region compared to cisgender men prior to receiving gender-affirming hormone therapy.

Contrary to several of the individual studies, no single-gene associations to GI were demonstrated for *AR*(CAG)n, *ESR2*(CA)n, *CYP19*(TTTA)n, and *CYP17* T>C. The results are also contrary to a previous meta-analysis that found significantly longer *AR*(CAG)n in transgender women compared to cisgender men (D’Andrea et al., 2020). Our *AR*(CAG)n analysis differs from D’Andrea *et al.,* (2020) in that we included data from Foreman *et al.,* (2019), and we obtained direct allelic information from the supplementary material of Fernández *et al*., (2018) instead of using mathematical estimation.

Assessment of the quality of these candidate gene studies revealed that most studies were either of low or moderate quality, with only two studies rated as high. A lack of representativeness of cases was a pervasive issue. All studies recruited transgender participants through gender clinics. Given that individuals who seek formal diagnosis/treatment of GI through gender clinics make up a small portion of a wider population who identify as transgender or otherwise gender-diverse (Arcelus et al., 2015; Meerwijk & Sevelius, 2017), this may contribute to selection bias. Some studies did not adequately control for ethnicity (Foreman et al., 2019; Lombardo et al., 2013). This is important, as genetic association studies require cases and controls to be ethnically homogenous to control for differences in allele frequencies and linkage disequilibrium between ancestral populations (Hellwege et al., 2017). Only three studies controlled for age of onset of GI and the sexual preferences of the participants (Fernández, Delgado-Zayas, et al., 2020; Fernández et al., 2016, 2018). It has been hypothesised that transgender individuals are not a homogenous group, but rather may be subtyped based on age of onset and sexual attraction (Blanchard, 1989), which may have support from neuroimaging studies (Guillamon et al., 2016). Based on this hypothesis, it may therefore be necessary to control for these factors in genetic studies to create as homogenous groups as possible, as different subtypes of gender incongruence may have unique developmental pathways involving different genetic factors.

Some limitations of this analysis should be acknowledged. Firstly, the power of the analysis was limited by the paucity of studies in this area. This is especially true of the analyses in transgender men, which only two studies reported data on. Secondly, the ability to interpret heterogeneity in the individual study results with reference to study characteristics such as ethnicity age of participants, clinical criteria used to define cases, age of onset of GI, or sexuality was limited due to the lack of studies. Thirdly, the combined data came from ethnically heterogeneous samples, making it difficult to determine how ethnicity may influence the results, due to potential variability in the allele frequency between ethnic populations (Zitzmann & Nieschlag, 2003). Lastly, we were unable to perform gene-gene interaction analysis due to the unavailability of genotype data.

It would be remiss to not address the ethical implications of conducting genetic research in transgender populations. Concerns have been raised that genetics research may negatively impact transgender people in several ways. For example, a survey of transgender or otherwise gender-diverse individuals found that people felt that genetics research could contribute to the gatekeeping of transgender identities (i.e., by encouraging people to think only those with a particular marker are ‘truly’ transgender), the gatekeeping of gender-affirming care (i.e., only those with a particular marker can access treatment), the pathologisation of transgender identities as mental disorders, the perpetuation of a binary conception of gender, or open the door to eugenic practices (Rajkovic et al., 2022). Although these are valid concerns, they should be weighed against the potential positives of conducting such research. Given the nature of complex traits, it is extremely unlikely that a single marker will ever be able to differentiate transgender and cisgender individuals; indeed, in this study, many cisgender men possessed the marker that was overrepresented in transgender women. On the contrary, demonstrating certain markers are overrepresented in people who experience GI could contribute to the de-stigmatisation of transgender identities in the eyes of the public, as several studies suggest that biological views of transgender identities are associated with positive attitudes towards transgender people, whilst the view that transgender identities are environmentally caused are associated with negative attitudes (Bowers & Whitley, 2020; Brown et al., 2017; Campbell et al., 2019; Ching & Chen, 2022; Elischberger et al., 2016, 2018; Greenburg & Gaia, 2019; Landén & Innala, 2000; Rad et al., 2019; Woodford et al., 2012). Additionally, studying the underlying genetics of gender identity may promote understanding transgender identities as representing natural human variation, rather than reinforcing a binary sense of gender. Such research may also have medical applications, such as for personalised medicine for transgender individuals. The genetics of sex hormones which may contribute to gender identity, as well as the use of exogeneous hormones for gender-affirmation, may affect how medical conditions are treated in transgender individuals (Moyer et al., 2019).

## Conclusion

This study is the largest genetic analysis of transgender individuals conducted to date. A significant association between a polymorphism in *ESR1* and gender incongruence amongst natal males was identified. However, future studies exploring genetic contributions to gender should employ genome-wide approaches. Candidate gene studies are prone to false-positives, likely due to the small effect sizes of genes that contribute to complex traits requiring large samples sizes to detect, and also due to ethnic heterogeneity between studies (Munafò et al., 2004). Further still, genome-wide association studies have revealed that our *a priori* assumptions as to what would constitute viable candidate genes are often wrong; significant associations in candidate gene studies do not replicate when the entire genome is scrutinised (Border et al., 2019; Hall et al., 2013; Johnson et al., 2017). Psychology is always highly multigenic (Plomin et al., 2016), and therefore no account of a genetic contribution to gender identity based on the examination of a few genes will be a complete one. So far, candidate gene studies have focussed exclusively on sex steroidogenesis genes, motivated by the hypothesis that the process of sexual differentiation of the brain is what is altered in transgender individuals. If this is the case, it is also possible that sex chromosome genes may have an influence, as brain differentiation has been shown to be influenced by sex chromosome genes before sex steroidogenesis begins (Corre et al., 2016). In light of recent evidence demonstrating the transgender individuals prior to receiving gender affirming hormone therapy show global DNA methylation differences to cisgender people (Ramirez et al., 2021), and that such therapy induces epigenetic changes that make their epigenetic profiles more similar to the opposite sex (Shepherd et al., 2022), it is possible that genes which control methylation in response to sex hormones could be implicated. Further still, it is possible that mechanisms outside of the sex differentiation pathway are involved in the development of GI which have not yet been considered.

## Supporting information

Meta-analysis data

## Declarations

### Availability of Data and Materials

The datasets used and/or analysed during the current study are available from the corresponding author on reasonable request.

### Code Availability

The R code used in this analysis are available from the corresponding author on reasonable request.

### Author Contributions

BA extracted data, conducted quality assessment of studies, conducted all statistical analysis, and wrote and edited the manuscript. VH conceived the project, extracted data, conducted quality assessment, and contributed to editing of the manuscript.

### Ethics Approval

Not applicable.

### Consent to Participate/Publish

Not applicable.

## Acknowledgements

We would like to thank Alice Anderson of Monash Health library for assisting in constructing the database search strategy.

## Compliance with Ethical Standards

### Research Involving Human Participants and/or Animals

Not applicable.

### Informed Consent

Not applicable.

